# One shot generalization in humans revealed through a drawing task

**DOI:** 10.1101/2021.05.31.446461

**Authors:** Henning Tiedemann, Yaniv Morgenstern, Filipp Schmidt, Roland W Fleming

## Abstract

Humans have the striking ability to learn and generalize new visual concepts from just a single exemplar. We suggest that when presented with a novel object, observers identify its significant features and infer a generative model of its shape, allowing them to mentally synthesize plausible variants. To test this, we showed participants abstract 2D shapes (‘Exemplars’) and asked them to draw new objects (‘Variations’) belonging to the same class. We show that this procedure created genuine novel categories. In line with our hypothesis, particular features of each Exemplar were preserved in its Variations and there was striking agreement between participants about *which* shape features were most distinctive. Also, we show that strategies to create Variations were strongly driven by part structure: new objects typically modified individual parts (e.g., limbs) of the Exemplar, often preserving part order, sometimes altering it. Together, our findings suggest that sophisticated internal generative models are key to how humans analyze and generalize from single exemplars.

## Introduction

Visual recognition and categorization of objects is vital for practically every visual task, from identifying food to locating potential predators. Humans can rapidly classify objects into familiar classes (e.g., Haxby et al., 2001, Thorpe et al., 1996, Mack & Palmeri, 2015, Fei-Fei et al., 2007) and form new classes when presented with sufficient example objects (e.g., Beeck & Baker, 2010; Gauthier, Williams, Tarr, & Tanaka, 1998). However, much less is known about how we infer new classes given just a small number or even only one example (“one-shot learning”; Feldman, 1992; Feldman, 1997; Ons & Wagemans, 2012; Richards, Feldman, & Jepson, 1992; Morgenstern, Schmidt, & Fleming, 2019), a skill acquired by humans at an early stage of visual development (Gelman & Markman, 1986; Gelman & Meyer, 2011; Gopnik & Sobel, 2000), with high consistency across observers (Feldman, 1992). From a computational point of view this ability seems virtually impossible: How can we predict the covariability of a category without having witnessed any variability? Given that any number of features within a category could potentially be variable or fixed, our ability to resolve generalization from sparse data must arise from certain strategies and biases. In contrast, machine learning approaches need upwards of thousands of pieces of data to successfully distinguish categories, showing both the difficulty of working with sparse data and the divide between machine learning and human approaches to categorization.

The state-of-the-art in psychology and computer science for understanding how observers generalize from few samples follows a generative approach. Specifically, it is assumed that observers have a deep understanding of objects, such that they know or can infer the elemental processes that generated the object. This potentially allows observers to identify behaviourally significant features, which in turn can be used to judge new samples (Fei-Fei, Fergus, & Perona, 2006; Feldman, 1992, 1995, 1997; Goodman, Tenenbaum, Feldman, & Griffiths, 2008; Goodman, Tenenbaum, Griffiths, & Feldman, 2008; Lake, Salakhutdinov, & Tenenbaum, 2015; Stuhlmüller, Tenenbaum, & Goodman, 2010). Consequently, understanding which features are identified as significant and how they are used to define a category is the key to this approach.

One possible strategy to differentiate between relevant and irrelevant features is to ascertain whether they could have emerged by chance. Random features do not tell us anything specific about the object at hand, on account of their random nature. Non-random features, however, will express themselves in a markedly more limited way, allowing us to gain usable insight into a categories’ feature-space. For example, the symmetrical arrangement of the four legs of a dog is very unlikely to occur by chance (i.e., if body and limbs were assembled at random). Thus, the arrangement of limbs would be a good candidate for a category-defining feature—whereas particular patterns of fur might appear more random and therefore would not be used for categorization (although it is possible that observers rely on some statistical summary for categorization, such as the regularity of the fur thickness or frequency). Previously, Feldman (1997) used the example of a dot on a line: New objects of that category are most likely to also be made up of a dot *on* a line as opposed to a dot separate from a line. The arrangement of the dot being on the line is deemed non-random and therefore, category-defining. To date, however, how we identify and interpret such category defining features and use them for generalization is a matter of debate (e.g., Serre, 2016).

A stumbling block in testing whether humans infer generative models of objects lies in current experimental methods in category learning. Typically, tasks exploring categorization and generalization from few samples ask observers to *discriminate* between presented objects (Ashby & Maddox, 2005; but see e.g. Stuhlmüller, Tenenbaum & Goodman, 2010), often varying along binary dimensions (e.g., thin vs. thick, black vs. white, square vs. circle; however, see Hegdé, Bart, & Kersten, 2008; Kromrey, Maestri, Hauffen, Bart, & Hegdé, 2010). However, this way of probing categorization or generalization results in responses that can be explained by a simpler and more efficient strategy than the generative approach outlined above. Specifically, observers could judge category membership based simply on similarity of statistical properties (e.g., area, perimeter, frequency, orientation) with no preference for any particular feature, and no ability to synthesize new variants. Another limitation of those tasks is that the definition of novel object classes through the selection of category features and their variability by the experimenter might differ from the features that independent observers might select to define categories.

To avoid these biases, and to tap into generative rather than discriminative processes, we used a generative one-shot task, in which observers were presented with single Exemplar objects and were asked to draw new objects (‘Variations’) from the same category, on a tablet computer. By analyzing the drawings relative to copies of Exemplars, and by asking other participants to (1) categorize them, (2) identify their distinctive features, and (3) compare them with Exemplar shapes, we can test whether the participants who drew the Variations truly generalized from single exemplars, and determine which features they focus on to do so.

This experimental approach allows rich new insights into how we select category-defining features and form categories. Specifically, by asking observers to *generate* objects rather than to *discriminate* between them, we gain direct access to observers’ generative models (Feldman, 1997), thereby tapping into our deep understanding of objects and their relevant features. Feldman (1997, 2000) also asked participants to draw a few new examples of basic visual concepts based on simplistic geometrical patterns (‘V’s, triangles and quadrilaterals). Here, by contrast, we used more complex Exemplar shapes varying in curvature characteristics and number of parts, and collected a large number of drawings, without discarding outliers. We reasoned that features that are reproduced across Variations are seen as significant features of the Exemplar shape. Similarly, other, random features that are not diagnostic of a category should not be reproduced, or, if reproduced, should not be seen by other observers as crucial to the object and its category membership. By asking other participants to compare, categorise and identify the significant features, we can work out *how* observers generalize and thereby describe their generative models.

## Results and Discussion

### Systematic generalisation in a generative one-shot categorization task (Experiments 1 and 2)

On each trial, one of the eight Exemplar shapes (**Fig 1a**) was presented in the upper half of a tablet computer’s screen. Participants were instructed to “draw a novel object that is not a copy of the Exemplar yet which belongs to the same category” using a digital pencil (**Experiment 1, Fig 1b**; see **Methods**). Exemplars were chosen to exhibit one or multiple prominent features. For each of the eight Exemplars, 17 participants (‘drawers’) each drew 12 Variations (yielding a total of 204 Variations per Exemplar). As a baseline, another group of participants (n = 15) were asked to copy the Exemplars three times as accurately as possible (**Experiment 2b**).

**Fig 1.**
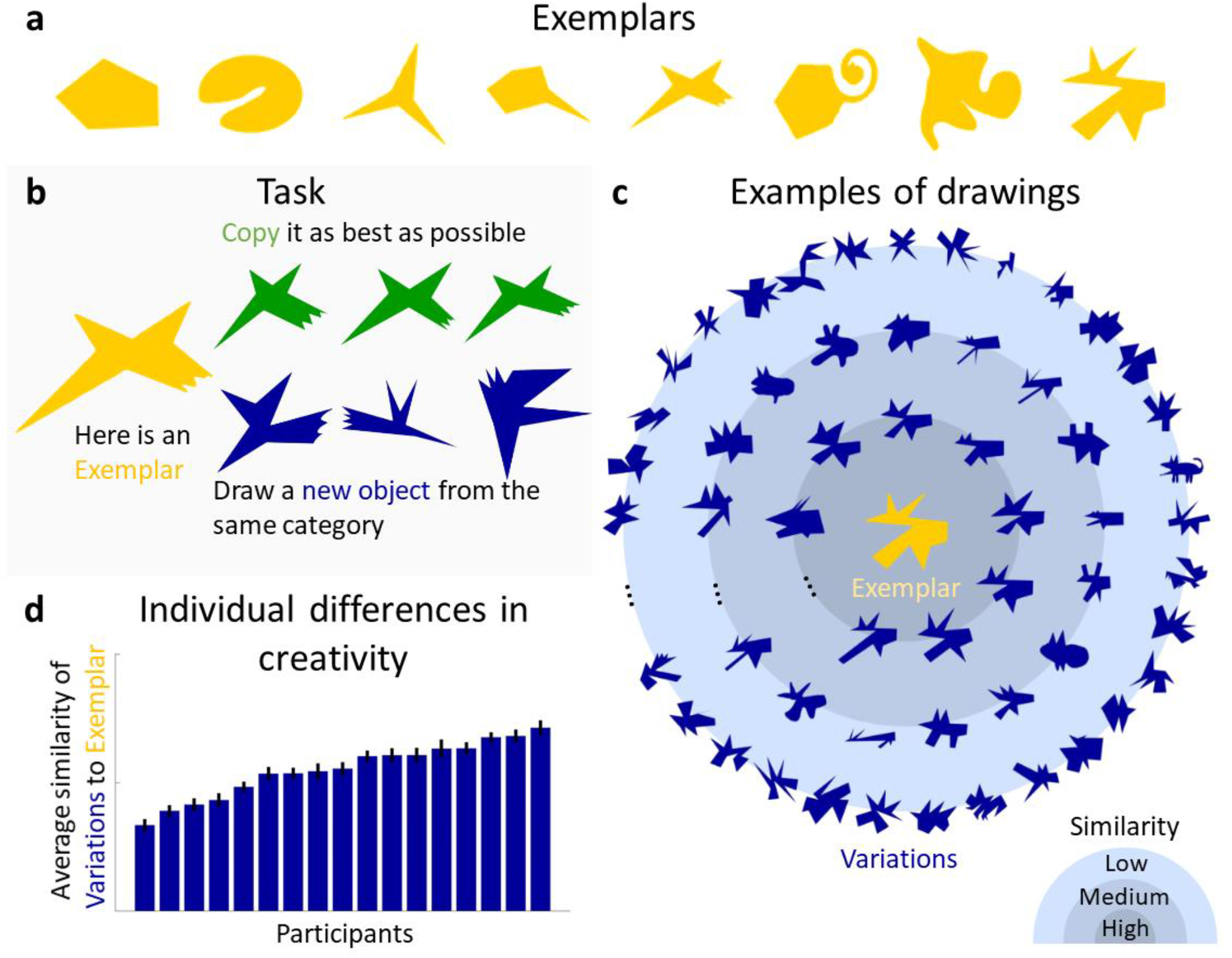
Generative one-shot categorization task and resulting data. **a**, Exemplars presented in the Experiment. **b**, in the task, a group of participants was presented with an Exemplar and asked to draw new objects from the same category. As a measure of baseline performance, another group of participants was asked to copy the Exemplar as accurately as possible. **c**, examples of the drawings (blue) generated in response to the shown Exemplar (yellow). Variations are plotted according to their perceptual similarity to the Exemplar, with more similar Variations closer to the center. **d**, Individual differences in creativity, defined by the mean similarity of their drawings to the Exemplar.

To assess the range and variety of the generated shapes, a new group of 12 participants rated the similarity of each Variation to the corresponding Exemplar (**Experiment 2**). **Fig 1c** shows examples of Variations for one Exemplar, with perceptually more similar drawings plotted closer to the Exemplar (see **Fig S1** for Variations of other Exemplars). The drawings illustrate the range of responses, with some shapes being very similar to the Exemplar while others differ considerably. **Fig 1d** shows the average similarity of produced Variations for each participant, demonstrating that participants varied substantially in how much their drawings deviated from the Exemplar, that is, in their creativity when producing Variations. To see whether the generated shapes were genuinely new objects or just slightly altered copies of the Exemplars, the previously drawn copies were compared to a subset of Variations using a 2-AFC task (**Experiment 2b**, see **Methods** for details), in which a new group of participants (n = 15) was asked to pick the shape that looked more like a copy of the Exemplar. In 95% of trials (chance = 50%) the copies were chosen, showing that the vast majority of a representative subset of Variations were perceived to be more different from the Exemplars than a mere copy.

### Responses represent distinct perceptual categories (Experiment 3)

We next tested whether the drawings represented distinct perceptual object categories or were merely random variations that did not form coherent groups. The similarity ratings for each Exemplar’s Variations from **Experiment 2** were split into subsets (40 bins) that spanned the full similarity continuum (see **Methods**). A new group of participants (n = 15) classified one Variation from each bin (for each Exemplar; 320 stimuli in total). In each trial, participants classified a randomly selected Variation into one of the eight Exemplars’ categories. The average percentage of correct classifications was high and well above chance; a binomial test indicated that the proportion of correct responses of 86% was significantly higher than chance level (12.5%,) (N = 4800 (number of judgements), K (correct responses) = 4111, p < 0.001)). **Fig 2a** shows the confusion matrix for true and perceived classes. In almost every instance observers were able to correctly choose the originating class of the Variations. Performing a one-sided binomial test on each cell shows that all cells except one (row 4, column 1) are significantly different from chance with all diagonal cells are above, all others below chance (Bonferroni-adjusted p-value of 0.015, adjusted by the 8 possible outcomes in each row). This demonstrates that there are no systematic mis-categorizations between particular Exemplar categories. Together, this is strong evidence that the Variations produced in the generative task are samples of robust perceptual categories. It suggests that drawers identified non-accidental features in the Exemplars and reproduced them in the Variations, allowing other observers to identify that they belonged to a common class. Thus, drawers effectively created distinct novel categories from just a single object.

**Fig 2.**
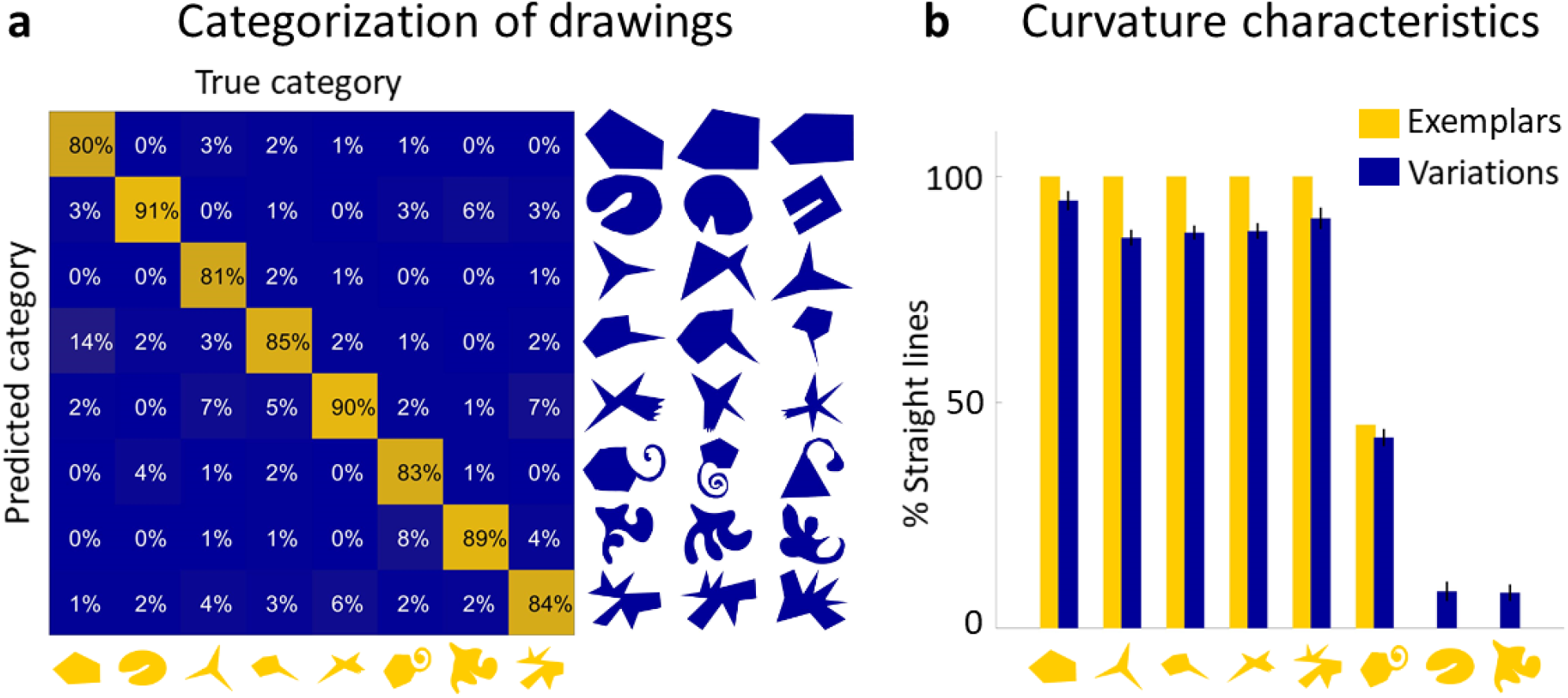
Drawings constitute a real perceptual category. **a**, the confusion matrix for true and predicted class shows that the vast majority of Variations were classified correctly. A subset of Variations that had to be categorized is shown on the right of the matrix with the Exemplars on the bottom. **b, Variations largely reproduce the global curvature characteristics of the Exemplar**. Curvature similarity across Variations and their Exemplars: Exemplars (yellow) ordered by the percentage of perimeter comprised of straight lines, together with average percentage of straight lines in all Variations of that category (blue).

### Identifying category-defining features

So far, our findings show that observers generated genuinely novel objects, which other observers could nevertheless assign to their corresponding category. This seems to suggest that drawers identified and reproduced those features of the object that are most significant and defining of the class. Thus, we next investigated *which* features were preserved, to which extent participants agreed about the most significant features, and the importance of these features for category assignment.

#### A. Global curvature features

One significant feature preserved in almost all Variations were the global curvature characteristics, namely whether the Exemplar consisted of straight or curved lines. We find that polygonal Exemplars tended to lead to polygonal Variations while curvaceous Exemplars led to curvaceous Variations (r = 0.99, p < 0.001; **Fig 2b**). This finding is broadly in line with the concept of non-random features that indicate generative processes (Feldman, 1992, 1997): A pencil tracing a random walk is unlikely to draw a straight line, so straight lines are considered evidence of a significant (i.e., non-random, class-defining) generative process, which are therefore preserved in Variations.

#### B. Identification and preservation of distinctive parts (Experiment 4)

Another noticeable feature of the generated shape-categories is that certain parts of the Exemplars seem to be more distinctive than others and remain so in Variations that share some or all of the part structure. We suggest that these distinctive parts are the main driving force for correct categorizations (**Experiment 3**). To address whether participants agree about which shape features are most distinctive, we showed a new group of participants (n = 10) the 8 Exemplars and 39 Variations from each of the categories, in random order. They were asked to paint up to three parts of each object’s silhouette, starting with the most distinctive part, followed by the second and third most distinctive parts. **Fig 3a** shows all responses for one shape together with the aggregated response, which indicates a high level of agreement across observers (see **Fig S2** for complete set). **Fig 3b** shows aggregated responses for further examples. Visual inspection suggests that for most categories, participants tend to consistently indicate specific parts as being the most distinctive, (e.g. indentation for category 2, spike for category 4, jagged feature for category 5, and twirl for category 6). Contrasting this data with randomized responses mimicking the human responses in terms of number of areas painted and lengths of consecutive areas painted, but with randomized placement of those areas (**see Methods** for details), shows significantly higher agreement among humans compared to chance: The mean consecutive area of a shapes’ silhouette with a high distinctiveness-score (above 75% of the highest possible score) for the human data made up 19% of a shape’s perimeter compared to < 1% for the randomized data (Two-sample Kolmogorov-Smirnov test, p < 0.001).

**Fig 3.**
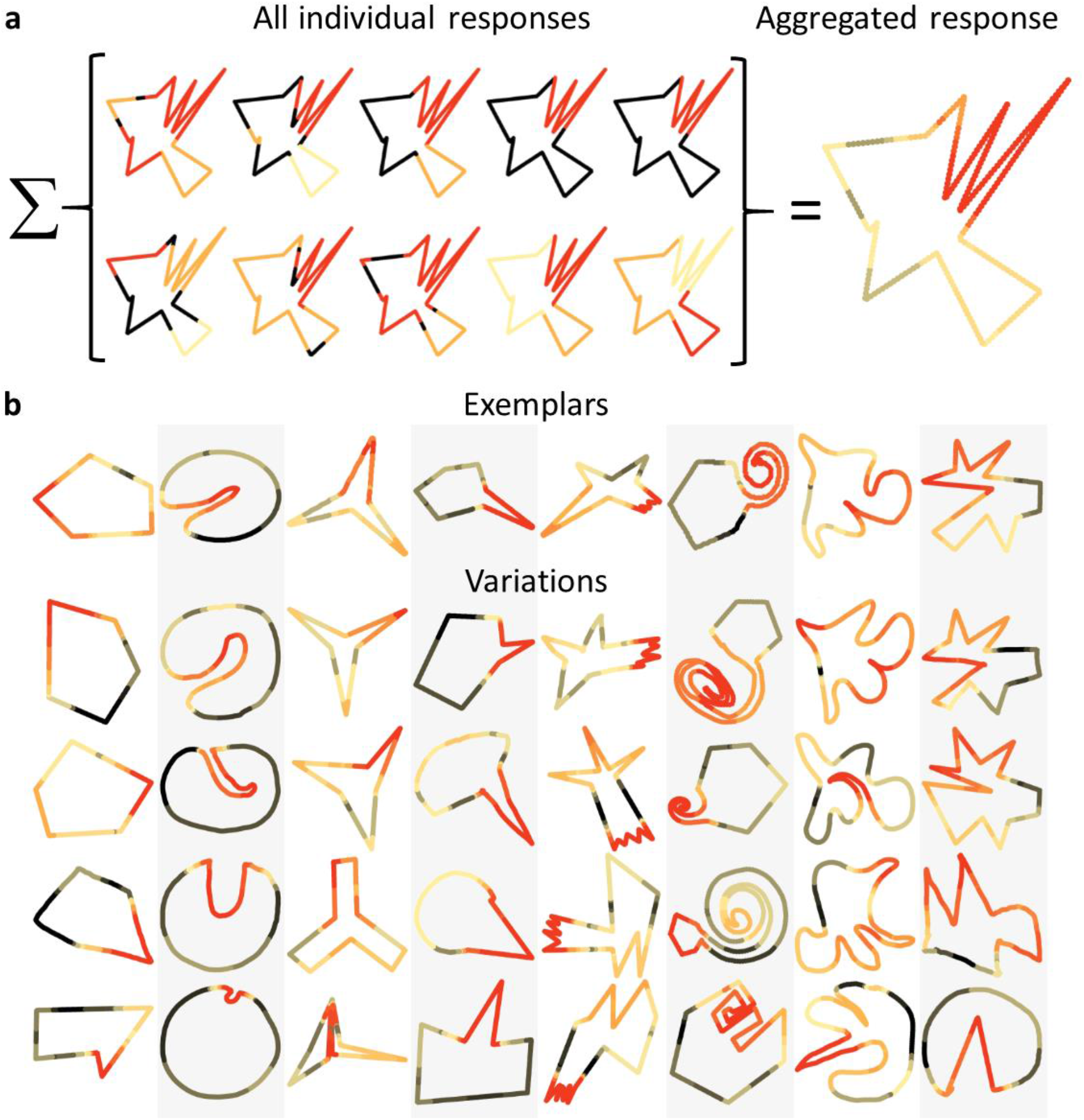
Observers agree on the most distinctive parts. **a**, individual responses of all participants for an example Variation, showing which parts were marked as most distinctive (red), second most distinctive (orange) and third most distinctive (yellow). Aggregating these responses results in the shape on the right. The redder each point on the contour, the more distinctive it was evaluated across all participants. **b**, comparing aggregated responses between Exemplars and some of the corresponding Variations suggests that in most categories similar parts were identified as distinctive across the Variations.

### Causal role of distinctive parts in categorisation (Experiment 5)

Finally, to confirm the significance of distinctive parts for category membership, we created new stimuli from Variations by replacing the most distinctive part—as determined from **Experiment 4**—with the most distinctive part from the Exemplar of another category. For comparison, we created stimuli where we replaced the same shapes’ least distinctive part with the least distinctive part of another categories’ Exemplar (**Fig 4a**; controlling for perimeter length—for more information on the creation process, see **Methods**). A new group of participants (n = 15) grouped these newly generated stimuli (280 with the distinctive, 280 with the indistinctive part swapped) with one of the Exemplars, similar to **Experiment 2. Fig 4b** compares the results with the percentage of correct categorizations in **Experiment 2**. Binomial tests indicate that the proportion of correct responses from **Experiment 2** (86%) was significantly higher than both swap-conditions (69% for indistinctive swap, p < 0.001, and 32% for distinctive swap, p < 0.001). Further, we find a large difference in effect sizes: while swapping an indistinctive part does have a small effect (Cohen’s *h* = 0.29), swapping a distinctive part has an immense effect on correct categorizations (Cohen’s *h* = 1.05).

**Fig 4.**
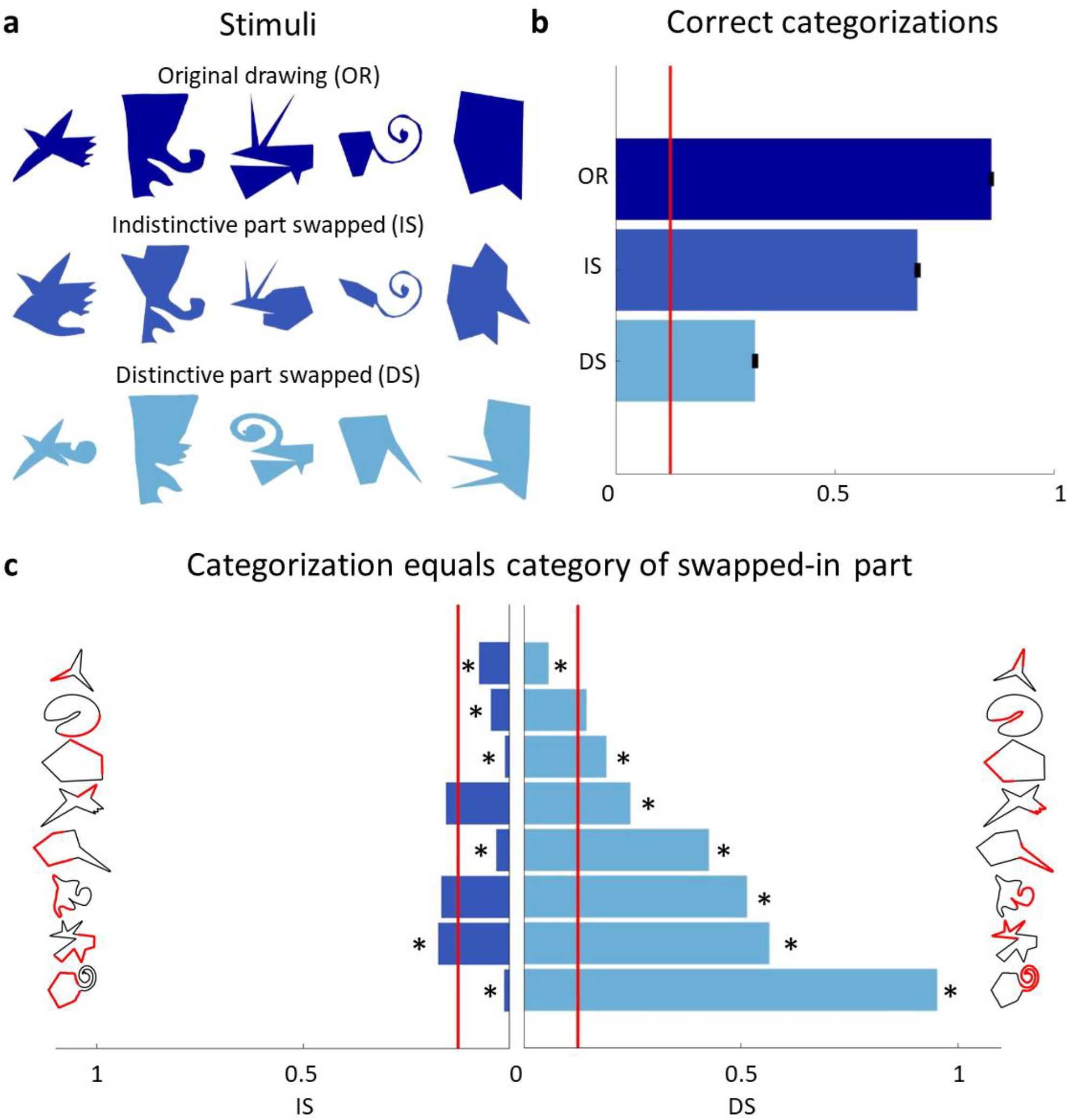
Distinctive parts are the main categorization cue. **a**, original drawings (top row) were altered so that either the least distinctive part was swapped with the least distinctive part of a different category (second row) or the most distinctive part was swapped with the most distinctive part of another category (third row). **b**, comparison of the percentage of correct categorizations for the original shapes (from **Experiment 2**), swapped indistinctive parts and swapped distinctive parts. **c**, bar plots showing how often the category of the swapped-in part determined the categorization choice. The indistinctive (left) and distinctive (right) parts are shown in red on the shapes’ silhouette. Bars significantly different from chance are marked with an asterisk.

This shows that distinctive parts played a driving force in categorization decisions, even though they often comprised only a small percentage of shapes’ contours. **Fig 4c** summarizes how often the category of the new, swapped-in part determined the categorization decision, separately for indistinctive and distinctive parts. Performing one-sided binomial tests on each of swapped-parts conditions (with a Bonferroni-adjusted p-level for 16 tests) shows that most distinctive parts strongly biased participant’s responses toward their categories, whereas only one indistinctive part had a significantly positive impact on it’s category-choice. Indeed, most indistinctive parts resulted in their category even being picked significantly below chance.

Previous studies showed that small to intermediate fragments of an image are sufficient for correct categorization (e.g. Hegdé, Bart, & Kersten, 2008; Ullman, Vidal-Naquet, & Sali, 2002). These informative fragments were defined by implicit analysis of the statistical distribution of features across the complete object category or categories (i.e., the fragments were learned during a training phase). Distinctive parts as identified in our experiments, in contrast, are identified from a single piece of data (i.e., one object) across observers and can therefore not be inferred from statistical feature distributions across many objects. Still, we suggest that observers may use statistical outliers *within shapes* to identify distinctive parts.

In summary, observers agreed on the most distinctive parts of shapes and use this information as one of the main cues for category membership. In line with this, when creating new shapes, these distinctive parts are reproduced somewhat modified but retaining the specific characteristics that make them both distinctive and signifiers of their category.

#### C. Part-related features (Experiment 6)

Considering the Variations in **Fig 1c**, it seems that many drawings approximately preserved the part-structure—i.e., arrangement and number of parts—of the Exemplars (this is especially salient when looking at the most similar Variations). At the same time, these parts were often modified in size, orientation or elongation. This signifies a potential strategy that participants might have used in the generative task: Starting from perceptual segmentation of the Exemplar into parts, they modify these parts (to varying degrees) and ‘put them back together’, either in the original or changed order.

To test whether participants used this strategy, another group of participants (n = 15) were shown one Exemplar at a time, paired with a Variation of either the same or different category, and asked to identify any corresponding parts between the two shapes. This allowed us to test whether part correspondence is stronger within categories than across categories—and how much of the Exemplar’s part-structure was retained in its Variations. If participants perceived parts as corresponding, they delineated each part by drawing a line, ‘cutting’ the shape into two parts on either side of the line, and then choosing one of the parts.

**Fig 5a** shows that correspondence was indeed significantly higher in same-category pairs than in different-category pairs. On average, 61% of an Exemplar’s area corresponded to some part of its Variations, with only 13% correspondence to Variations of other categories (**Fig S3a**)—showing that a substantial portion of the Exemplar’s parts were preserved in its Variations. **Fig 5b** shows aggregated mappings of correspondence for two of the categories, with the Exemplar on the left and rows of corresponding Variations in descending similarity to the Exemplar (see **Fig S3b** for complete set). Each point of the Variations’ contour is colored the same as the Exemplar’s contour to which it was perceived to correspond most. If a point was perceived to correspond to a whole section of the Exemplar (e.g. the green ‘nose’ of the Exemplar in the bottom row of **Fig 5b**), then it was colored the same as the circular median point of that section. Color saturation indicates how often correspondence was seen for each point, with higher saturation indicating stronger correspondence. This visualization shows the generally high level of agreement about corresponding parts between participants. Overall, these findings demonstrate that Exemplars and associated Variations share a considerable portion of their parts, indicating that drawers preserve identifiable parts in their drawings even though specific geometrical properties of the parts varied.

**Fig 5.**
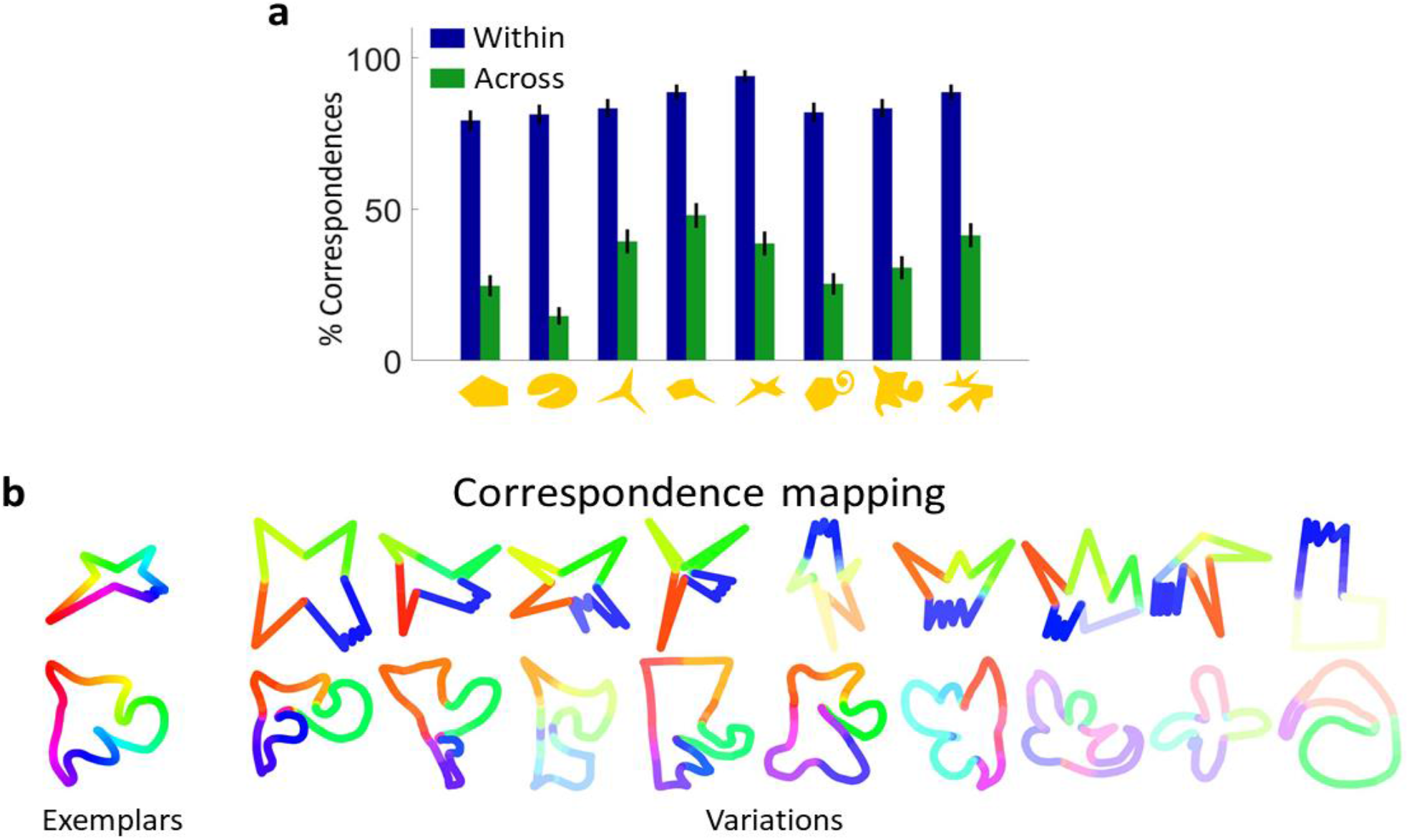
Variations and Exemplars of the same category share most of their parts. **a**, comparison of percentage of correspondences seen within (blue) and across (green) categories, showing that significantly more correspondence was perceived within categories. **b**, aggregated mapping of correspondences for two categories, showing high agreement between participants about corresponding parts.

Cross-referencing this data with distinctiveness scores from **Experiment 4** allows us to test whether distinctive parts within a category are indeed conserved (i.e., whether the most distinctive parts in each shape are seen to correspond). For each corresponding part-pair, average distinctiveness scores of both parts were calculated. Comparing the scores of all Exemplar parts with the scores of all Variation parts of the same category shows a high correlation (r = 0.63; p < 0.001). This suggests that distinctive parts remain distinctive even when modified or shuffled within the overall part-structure. Equivalently, indistinctive parts in Exemplars tend to remain indistinctive in Variations, lending further support to the finding that participants modify individual parts and rearrange them to create new shapes.

### General Discussion

One-shot categorization is a formally under-constrained inference problem (Feldman, 1992). Given only a single exemplar of a category, there is an infinite number of sets containing that exemplar, any of which could in principle be the true category from which the exemplar was drawn. It is thus remarkable that humans seem to be able to draw consistent conclusions about category membership from such sparse data (Feldman, 1992, 1997). Their judgments presumably reflect assumptions about how—and by how much—objects within a category tend to differ from one another, which constrains the space of variants that are deemed likely. Yet how this occurs remains elusive.

Most experimental studies in the field of categorization use sets of pre-selected stimuli to define categories and discriminative tasks to test their hypotheses. The main drawback of this approach is that the assumed category-defining features are determined by the selected stimuli and might therefore differ from the features used in unrestricted categorization decisions. Moreover, discriminative tasks allow for simple strategies based on comparisons between objects, rather than probing the visual system’s internal generative models of objects which are thought to be central to how humans and machines can learn categories from sparse data (e.g. Stuhlmüller, Tenenbaum & Goodman, 2010).

Here, by contrast, we use a generative one-shot categorization task to tap directly into generative models and human creativity—and to identify category-defining features by allowing participants to generate their own new category samples, rather than merely selecting between experimenter-generated alternatives. The resulting Variations are thus shaped by the features that participants consider important in the context of their previous experiences and by internal visual and imagery processes. In principle, different observers might consider different features as significant so that across observers no coherent object categories would emerge. However, in our findings this was clearly not the case with a high degree of agreement between observers, suggesting general principles of how humans analyze single objects and extrapolate new category-members from its features. Participants created Variations for different Exemplars varying in curvature characteristics and number of parts, resulting in a large data-set of over 1600 drawings, which were significantly more variable than mere copies of the Exemplars. Yet, despite the wide variety of Variations, the overwhelming majority of a representative subset of these shapes was correctly grouped with the originating Exemplar by independent observers, showing that our task created real perceptual categories.

A key finding is that one of the most important strategies to create new shapes was part-based. It seems that to create a new shape, the Exemplar was first segmented into its perceptual parts. Then, these parts were modified and recombined to form a new shape. Sometimes the original part-structure was preserved, resulting in a shape looking like a globally transformed (e.g., warped) version of the Exemplar. At times the parts were shuffled, resulting in a shape with a different part ordering. Sometimes only a subset of the original parts was used, or parts were added with respect to the Exemplar. **Fig 6** visualizes these proposed steps. However, even when the part-structure was altered substantially, other observers were able to identify corresponding parts between Variations and Exemplar, with a high degree of consistency. Evidently, drawers based their Variations on assumptions about how much a part—or the shape as a whole—could be varied while still retaining its identity. Importantly, these assumptions seem to be shared by other observers. In addition to the most prominent strategy outlined above, a range of other, more complex strategies were also used to generate shapes that grouped with the Exemplar, and even some that did not (**Fig 7**).

**Fig 6.**
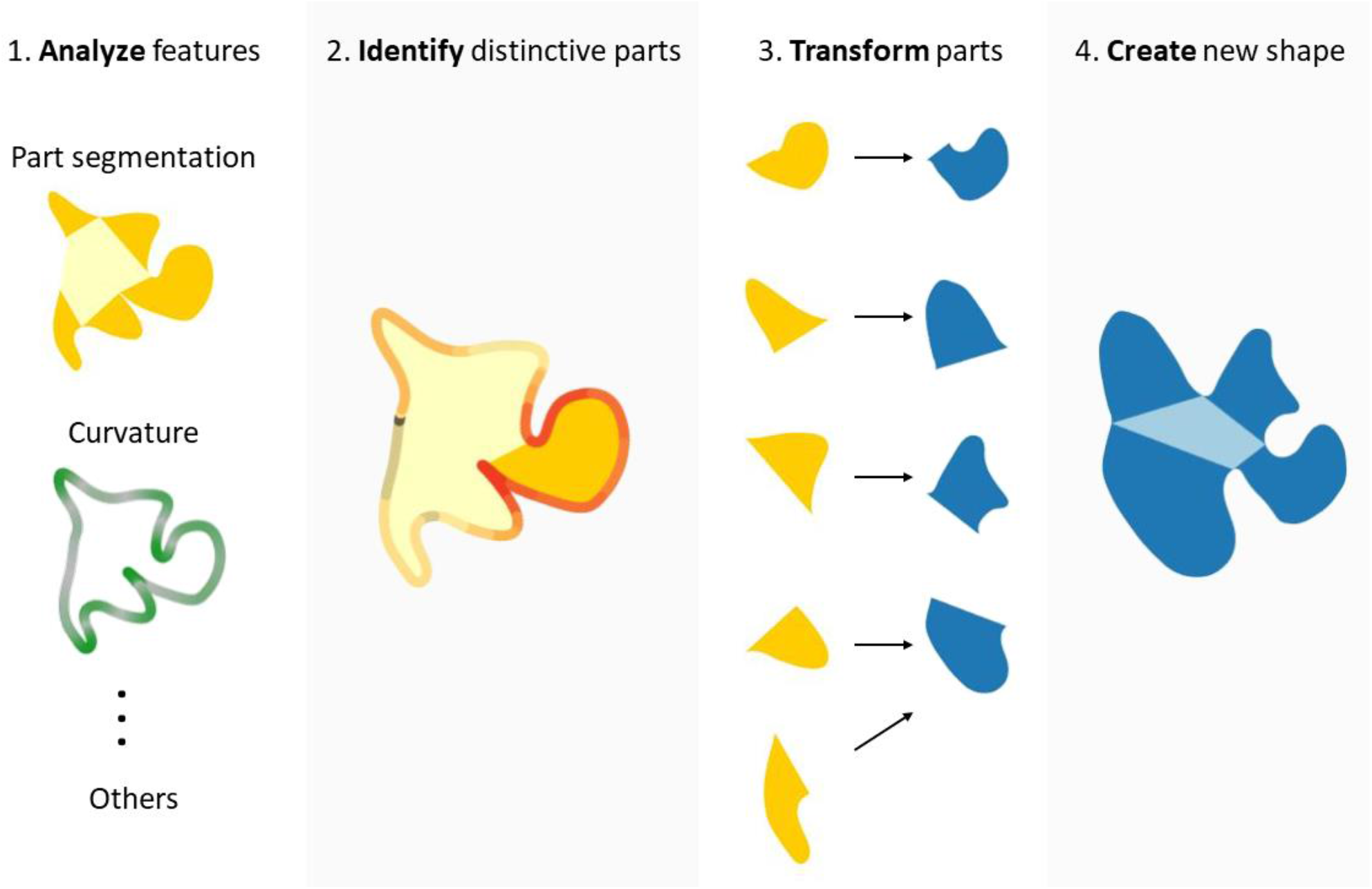
Proposed steps of shape creation. After analyzing shape features and identifying distinctive parts, the individual parts get transformed to form a new shape. In step 3 parts can get added, merged or removed and the order of parts might be changed.

**Fig 7.**
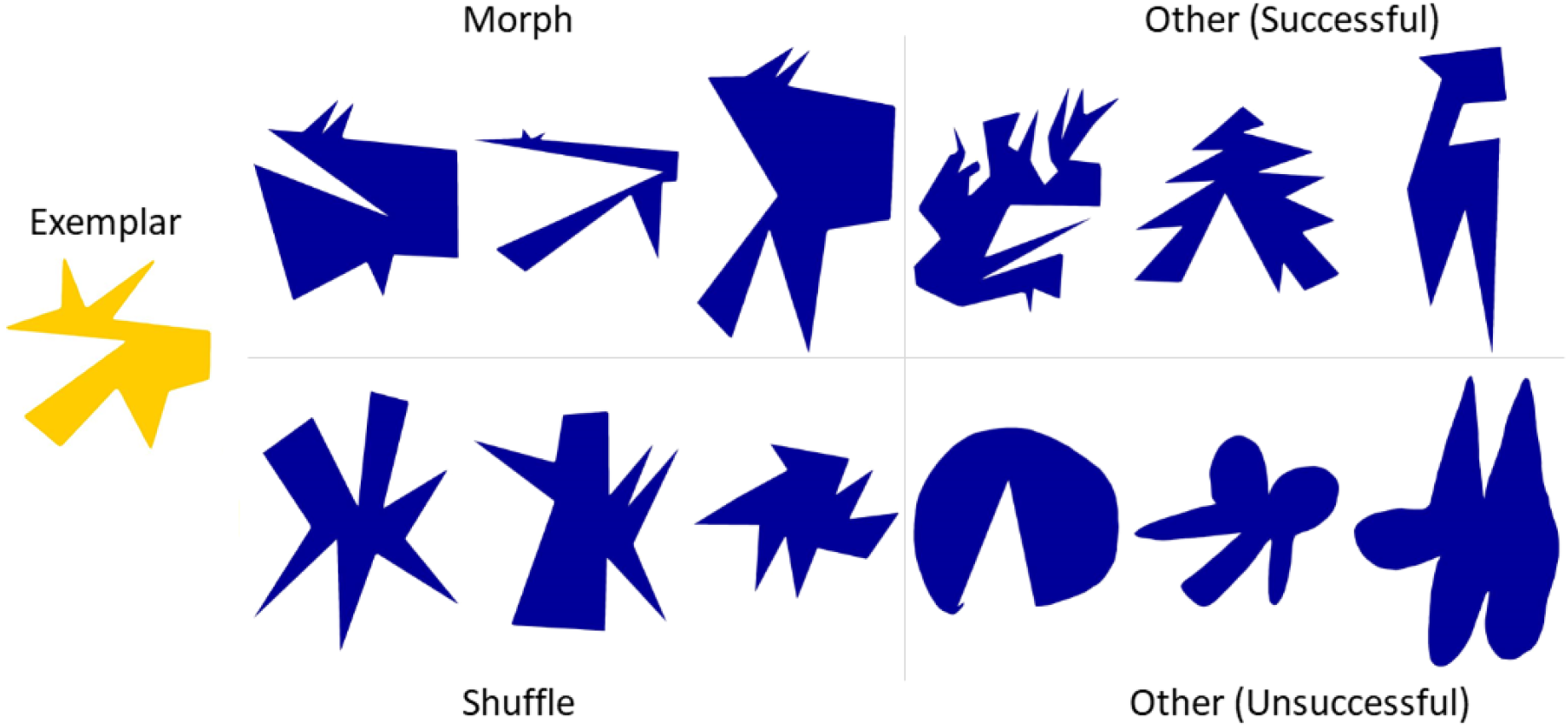
Different strategies used by participants. Morph, shuffle and other strategies, both successful (i.e., shapes were correctly classified almost all the time) and unsuccessful strategies (i.e., correctly classified far below average).

Many of the generated shapes not only shared most of their parts, but, strikingly, specific parts were also reliably perceived as more distinctive than others and were the main catalyst driving categorization-decisions, even if they comprised only a minority of the shapes’ area. This showcases a strength of the generative one-shot categorization task: Category-defining features were extracted, reproduced in modified form and recognized by other observers across a range of different tasks. As a consequence, we can be highly confident with respect to the features that drive categorization-decisions, without relying on presumptions about likely features and their variability. This suggests that a crucial stage in one-shot categorization is the identification of those features within an object that are most likely to be ‘distinctive’ for defining the category.

What is the basis of such inferences? Given only a single exemplar, how are the ‘most distinctive’ features determined? We suggest that observers’ decisions are related to processes that identify signatures of the underlying *generative processes* responsible for creating the observed shape and which thus define the category. This is closely related to Feldman’s (1992, 1997) theory of non-accidental features—statistical relations that are unlikely to occur under a random model (e.g., collinearity of features) are evidence of the operation of non-random generative processes. Similarly, parts that have statistically distinct properties from the rest of the shape—or which are statistically distinct from parts of objects seen previously—are likely evidence of a category-defining process. Consistent with this idea, we find that parts that were deemed distinctive were also those that had the most significant effect of categorization (e.g., the ‘twirl’ feature in category 6). On a more global scale, curvature characteristics (straight or curved lines) of Exemplars were preserved almost perfectly in most drawings, showing that these features were also deemed non-random. Conversely, random features were altered more freely: Since the presented shapes were abstract, the part-arrangement held no meaning and was therefore free to be modified (in contrast to familiar objects like a chair or a human), resulting in Variations with changed part-ordering from the Exemplar. Yet an important open question is which geometrical properties of parts are used to determine their status as outliers, and how these properties are themselves determined, based on familiar objects and innate processing constraints in the visual system. Another important open question is how parts are parameterised so that identity-preserving variations can be generated. Skeletal representations (Feldman & Singh, 2006) offer a promising avenue for potential representations.

Taken together, the results of our experiments suggest that humans are not merely passive observers, who assign objects to categories based on their relative similarities in a fixed feature-space. Instead, a key aspect of human generalization is our ability to identify important signatures of generative processes and then to run internal routines that actively synthesise new objects that we have never actually observed. Thus, although drawing tasks present experimenters with challenges—particularly in terms of analyzing the resulting drawings—they also provide an exciting avenue into research not only about object categorization but also human creativity. As seen in **Fig 1d** there is a notable difference between participants as to how much they tended to deviate from the Exemplars, provoking the question just what drives these individual differences. Another particularly fascinating open question is the extent to which the synthesis of novel objects by observers recruits physical simulation processes (Battaglia et al., 2013). It seems plausible that experience with the natural ways that objects, materials, animals and plants move and change shape over time (Schmidt & Fleming, 2016; Schmidt, Spröte & Fleming, 2016) might influence the types of variations that we tend to imagine (e.g., articulating limbs into different poses). The fact that observers can derive diverse but constrained variations from a single exemplar suggests a deep perceptual understanding about the ways things in the natural world tend to vary.

## Methods

Participants were recruited through university mailing lists. All participants gave informed consent before the experiments. **Experiment 1** was conducted on a touchpad computer, all others on a computer.

### Experiment 1: Generative one-shot categorization task

17 participants drew 12 new shapes for each of the 8 Exemplars (overall 1632 shapes). They were instructed to draw a new object, belonging to the Exemplar’s class, that is not just a mere copy of the Exemplar. The Exemplars were shown in randomized order in the upper part of an iPad Pro (12.9”) oriented in portrait mode and observers’ drawings were on the lower part of the screen. While drawing, participants could switch between drawing freely and drawing straight lines. Participants could clear the current drawing or undo the last drawn segment. After putting down the pen they could only continue drawing from the end-point of the last segment to prevent gaps between lines. After finishing a shape, the drawing area was cleared and a new object could be drawn. A shape could only be finished if the contour was closed, i.e. the first and last point drawn were on the same spot. Participants were instructed not to draw any overlapping or crossing lines.

### Experiment 2: Perceived similarity to Exemplars

For each category, 12 new participants judged the perceived similarity of the drawings to their originating Exemplar. First, all Variations of a category were shown in grids of 3 by 4 shapes consecutively to give the observers an idea of the range of shapes to be judged. In total, participants viewed 17 grids and 12 shapes. They were then presented with the Exemplar on the far left of the screen. On the screen bottom, participants used the mouse to drag and drop 3 randomly chosen Variations from that category into the upper part of the screen based on similarity. The closer the shape was placed on the x-axis towards the Exemplar, the more similar it was judged to be. Participants were instructed to try and keep a consistent scale of similarity, meaning that equally similar shapes relative to the Exemplar should be placed in the same area of the screen, regardless of what trial the shapes were presented or how similar those shapes were to each other. After placing 3 shapes relative to the Exemplar, a button press revealed the next 3 shapes on the screen’s bottom portion. Already placed shapes were greyed out but could still be adjusted in position if so desired. To prevent the screen from getting cluttered, old shapes disappeared after a few trials so that only 12 shapes were shown at one time. For each of the participants responses, the similarity values were normalized between 0 (the Exemplar) and 1 (the least similar drawing). The final similarity value of each drawing was averaged over these normalized responses.

### Picking stimuli from similarity space

In multiple experiments, we selected a subset of shapes from each category ensuring that subset consisted of the whole range of similarities. As an example, a subset of 20 shapes was created by taking all shapes of a category (minus the Exemplar) and dividing them into 20 equally sized bins spanning the similarity space. For each bin, we selected the shape with the lowest variance in similarity judgements. If a bin was empty, we searched neighbouring bins for a shape not yet used.

### Experiment 2b: Comparing copies and new drawings

15 participants were instructed to copy each Exemplar three times as best as possible with the drawing interface used in **Experiment 1**, resulting in 45 copies of each Exemplar. Then 15 new participants were shown one of these copies, along with the Exemplar and one Variation of that category per trial. The task was to pick the shape that was a copy of the Exemplar. The 45 Variations per category were chosen in the aforementioned way and randomly paired with one of the copies. Each category comprised one block of trials with blocks being randomized in order, resulting in 45 * 8 = 360 trials. These trials were repeated three times resulting in 1080 trials overall. In each repeat the order of blocks and pairings of copies and Variations were randomized.

### Experiment 3: Perceptual category-membership experiment

15 participants sequentially judged which Exemplar category 40 drawings (chosen as described in **Picking stimuli from similarity space**) belonged to. If participants were unsure about the category-membership they were instructed to pick the category they thought the shape belonged to the most.

### Experiment 4: Marking distinctive parts experiment

10 participants were sequentially shown a subset of 39 shapes (chosen as described in **Picking stimuli from similarity space**) for each category plus the Exemplar. They were asked to mark the most distinctive areas of that shape. They could freely paint on the shape’s silhouette and were not constrained to paint only consecutive parts. After painting the most distinctive part or parts in red they could switch to the next lower distinctiveness-tier (orange) and paint the second most distinctive areas. After that they could switch to yellow for the third most distinctive area. At least one part had to be painted red, the lower distinctiveness-tiers were optional.

To aggregate these responses (as in **Fig 4a** and **S2**), each point of the contour was given a score, with each red response adding 3, each orange 2 and each yellow adding 1 to the score of that point. These scores were then normalized between 0 (score = 0) and 100 (highest possible score).

### Creation of randomized responses for Experiment 4

For each categories’ responses, we computed the percentage of painted perimeter of each distinctiveness-tier. In addition, we computed the distribution of lengths of each tier’s consecutive painted areas and the distribution of number of non-consecutive areas per tier and category. With these distributions 10 (the number of participants in **Experiment 4**) randomized responses were created for each of the shapes from **Experiment 4**, meaning the average number of areas and consecutive lengths mimicked the human distribution, while the placement on the shape was random.

### Swapping distinctive parts stimuli generation

First, we segmented each shape of **Experiment 4** into one distinctive and one indistinctive part. The distinctive part was the largest consecutive part of the shape with an aggregated distinctiveness score higher than 75 in each point. The indistinctive part was the rest of the shape.

In “distinctive part swapped” shapes, we swapped the distinctive part with a distinctive part of another categories’ Exemplar. In “indistinctive part swapped” shapes, we swapped an indistinctive part of the shape with the same number of points as the distinctive part that had the lowest average distinctiveness score. Since points were equally spaced along the contour both parts had the same perimeter-length. In either case the swapped-in part was rotated and either compressed or stretched to fit the gap as best as possible.

In each case a drawing from the subset from **Experiment 4** (excluding the Exemplars) was used as the base shape. In this way for each of the 8 categories, 5 shapes were created with parts swapped from one of each of the other 7 categories both once with the distinctive part and once with the indistinctive part swapped, resulting in 8 * 7 * 2 * 5 = 560 shapes.

Because this process sometimes created shapes with large self-intersections, the final 5 shapes for each condition were hand-picked to form well-formed, artefact-free shapes.

### Experiment 5: Swapping distinctive parts experiment

15 participants conducted a repeat of the “Perceptual category-membership” experiment with 560 stimuli generated as explained in the section **Swapping distinctive parts stimuli generation**.

### Experiment 6: Corresponding parts experiment

15 participants participated in this experiment. For each Exemplar, a subset of 10 drawings was chosen in the aforementioned way. Participants were shown an Exemplar on the left and a drawing on the right, either belonging to the same or a different category. They were then asked if they saw any corresponding parts between these shapes. If not, the next shape-pair was shown. Otherwise they were asked to pick the corresponding parts by first picking the part in the Exemplar and then the corresponding part in the drawing. To pick a part, the two delineating points of the part were to be clicked in succession, creating a line between these points within the shape. After that, the polygon on either side of the line could be chosen as the final part. A part could only be picked if the resulting line between the points did not intersect with other parts of the shape. Instead of picking a part, the rest of the shape (comprised of anything not yet picked) could also be picked, indicated by a red dot at the centroid of the remaining shape. Any section already picked could not be used for another part. Any number of part-pairs could be picked, unless the remaining shape was picked, after which the trial was ended, since no more unpicked parts remained.

Each Exemplar was paired with 10 corresponding and 10 Variations of another randomly chosen category, resulting in 8 * 20 = 160 trials.

### Perceived Exemplar complexity

20 participants arranged the 8 Exemplars in order of perceived complexity from simple to complex. The average rank of these responses was used to order the Exemplars and their categories in most plots of this paper.

## Acknowledgements

Research funded by the Deutsche Forschungsgemeinschaft (DFG, German Research Foundation)–project number 222641018–SFB/TRR 135 TP C1), and by the European Research Council (ERC) Consolidator Award “SHAPE”–project number ERC-2015-CoG-682859.

## Supplemental figures

**Fig S1.**
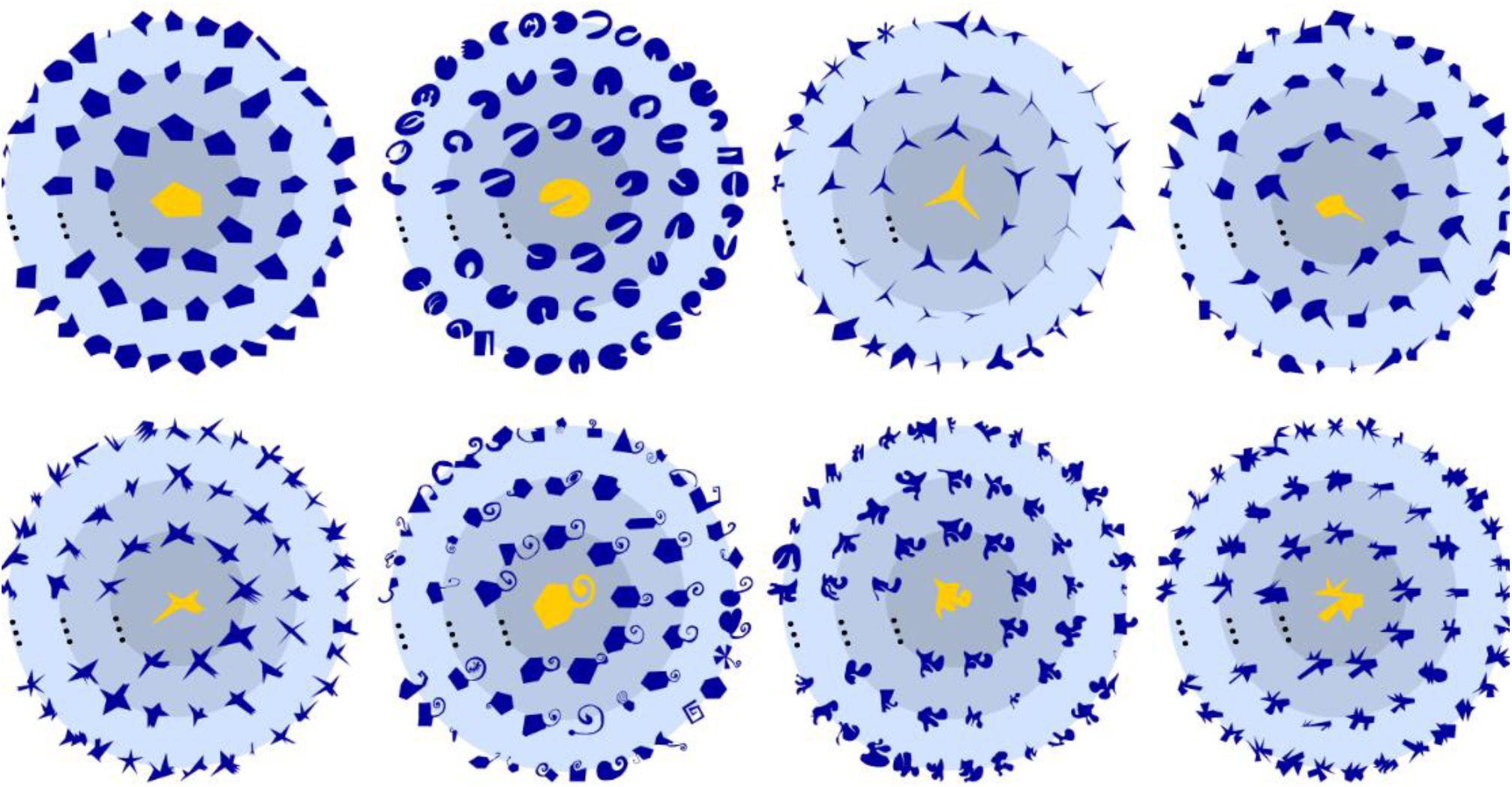
A Subset of Variations (dark blue) created for each Exemplar (yellow, center). Variations closer to the center were deemed more similar to the respective Exemplar.

**Fig S2.**
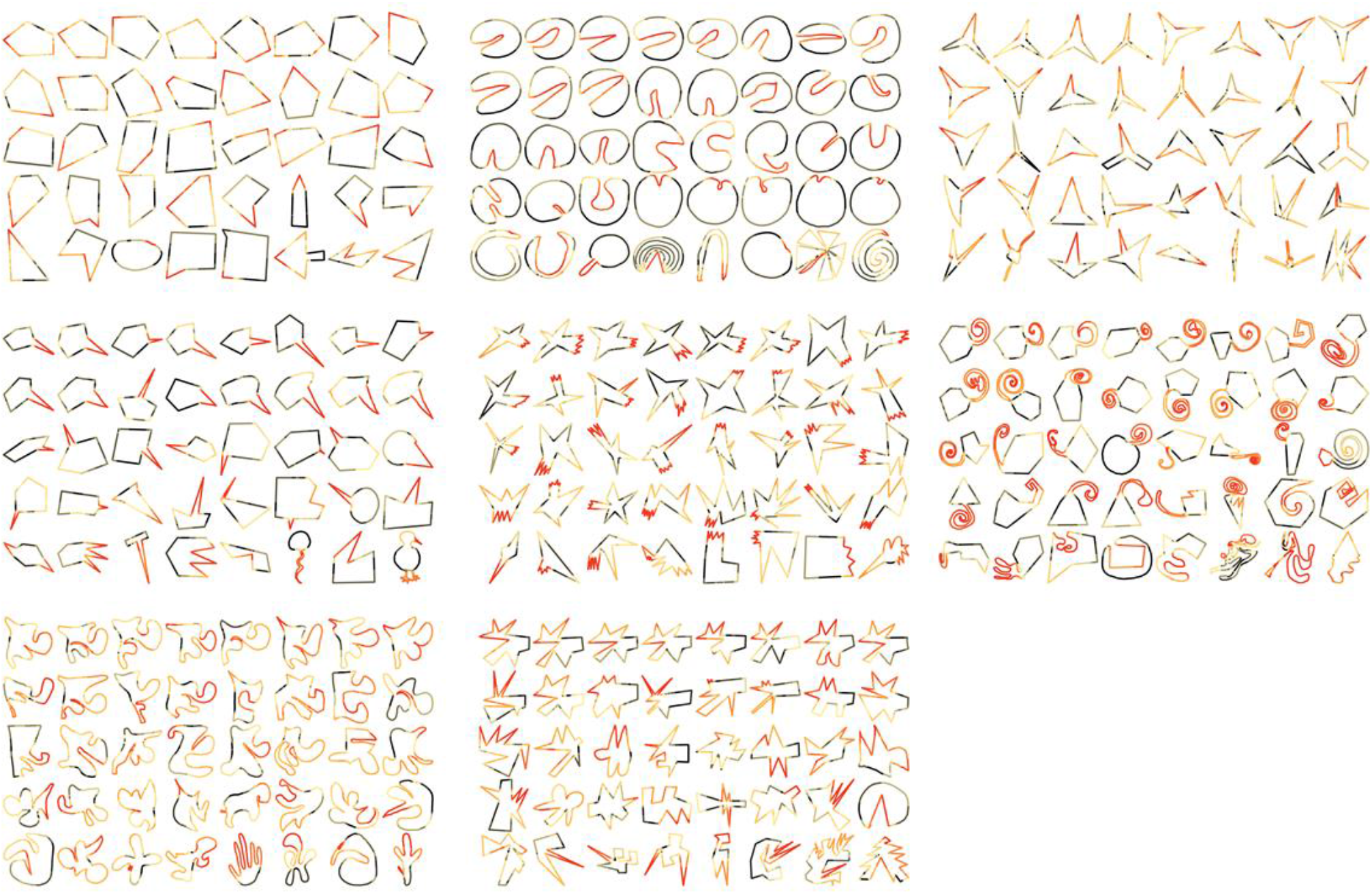
Aggregated distinctiveness scores for all shapes. The first shape in each category is the Exemplar, with the following shapes representing all other shapes used in **Experiment 4** in descending order of similarity to the respective Exemplar. Distinctiveness is represented by color of each point from black (completely indistinctive) through yellow and orange to red (very distinctive).

**Fig S3.**
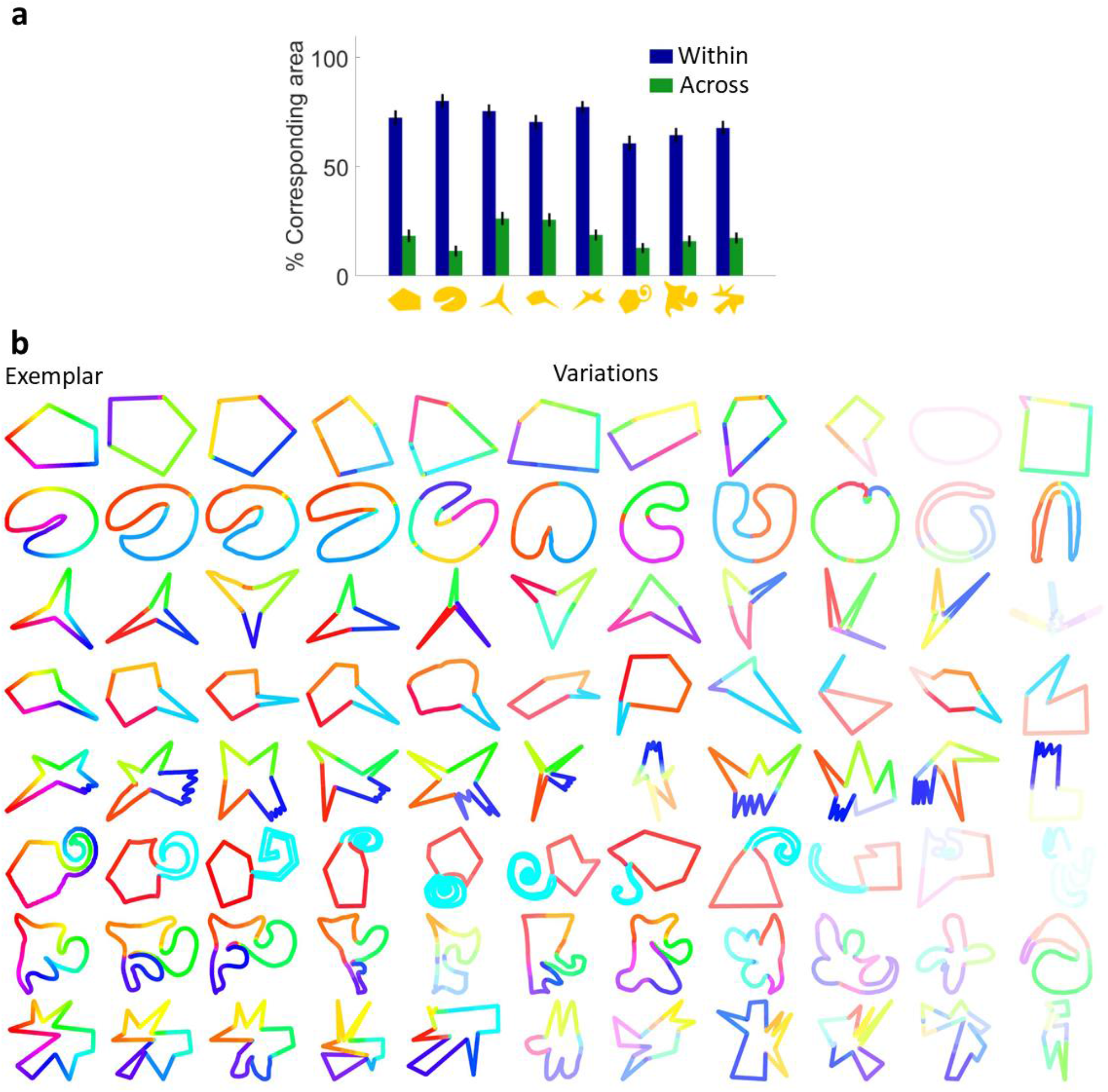
Correspondence data for whole subset. **a**, Mean corresponding area seen to correspond for each category’s Exemplar with Variations from other categories (green bars) and from the same category (blue), showing both the large difference in area corresponding within and across categories and that on average a large majority of each within-category shape pair was seen to correspond. **b**, correspondence mapping for each Exemplar and Variation used in **Experiment 6**.

